# Interspecific continuity of normative systems (the end of the paradigm of human exceptionalism)

**DOI:** 10.1101/2023.08.19.553974

**Authors:** L. Gandarillas Chicote, P.M. De La Cuesta Aguado

**Affiliations:** Seminario de Derecho Penal, Facultad de Derecho, Universidad de Cantabria, avda. de los Castros s/n, 39005, Santander, Spain; Cabárceno Nature Park, 39690, Obregón, Cantabria, Spain

## Abstract

After fifteen years observing a troop of western lowland gorillas (*Gorilla gorilla*) outside their natural habitat under human care we believe we can demonstrate that gorillas govern their behavior and social interactions by norms functionally equivalent to legal norms that pursue group stability and social peace maintenance and can be “translated” into concepts created for legal science. Consequently, we conclude that also in the legal sciences and, especially, in the juridicial-criminal science, the paradigm of human exceptionality must be overcome to open the doors to reflect on the possibility of establishing the paradigm of continuity between human and non-human normative systems.

## Main Text

The social sciences respect the paradigm of human exceptionalism –of the essential difference between human nature and animal nature– even when evolution is scientifically proven. Law is a product of human society, however, our observations invite us to conclude that also in the legal sciences, and especially in legal-criminal science, the paradigm of exceptionality must be overcome to open the doors to reflect on the possibility that similarities between species transcend the biological and are projected through behaviour, so that regulatory models with common elements can also be identified.

Here we show how a troop of western lowland gorillas in captivity is governed by norms functionally equivalent to legal norms and how these can be “translated” into concepts created for legal science. This not only advances the understanding of the ethology of gorillas but also the functions of criminal law and punishment in our societies.

Legal norms establish patterns of behaviour with a claim of generality and stability^1^ and sanctions for its infringement. They are intended to motivate action^2^ and are coercively imposed by an external agent^1^. A legal system is an orderly set of (legal) norms based on values with rules that regulate its internal functioning. Human society is built with law and because of law (*ubi societas, ubi ius*)^3^ and the question we raise is whether this is also the case in non-human societies.

A society is conformed by a set of individuals and interrelated social situations that form a heterogeneous set with the capacity to coordinate beyond the individual actions and particular intentions of each member^4^ and it is not exclusive to the human species^5^. The type of society that can emanate from each species depends on the characteristics of its habitat and its social development^6^. For a society to be constituted, the individuals who make it up must possess minimum capacities: a communication system^7^, the ability to carry out prosocial behaviours^8,9^, a sense of social regularity or justice^10,11,12^ that manifests itself in behaviours such as respect for possession^13^ or police intervention^14^ and, in addition, they must possess culture, defined as the transmission of information between individuals that is not inherited through genes^15,16^. These abilities are not unique to humans and are also found in gorillas, who can convey emotions, intentions, environmental characteristics, greetings, and even reach agreements through their communication system that uses vocalizations^17,18,19^, gestures^20^, and even body odour^21^. They also show prosocial behaviours^22^, aversion to inequality^23^ and police behaviour^24^, perform cultural transfers of techniques^25^ and behaviours^26^ and present a cultural flexibility dependent on the environment^27,28^.

Our research is aimed at studying a troop of western lowland gorillas at Cabárceno Nature Park (Spain), which began to establish on April 27, 2007 with a pair of adult western lowland gorillas (*Gorilla gorilla gorilla*), both born in the wild in Equatorial Guinea. Later, other females of different origins were added to the group until they formed the current family that consists of a silverback male, 4 adult females and 3 offspring born in Cabárceno (F1). The keepers are trained to ensure that the animals can maintain their natural behaviours. Although this troop of gorillas lives in captivity, it maintains the basic structures and behaviours of a social group in their natural habitat, which allows for continuous observation and research of their behaviour.

The research has been developed in two phases. A first phase of observations *ad libitum* over 6813 hours (F1) between 2007 and 2021, from which we have created a catalogue of behaviors (F2) whose results we attach in Figure F3. Secondly, we have developed another phase of systematic research using ethograms for 21 hours (F4, F5). In both we have analyzed the behaviour observed applying the knowledge and concepts of criminal science.

**Fig. F1:**
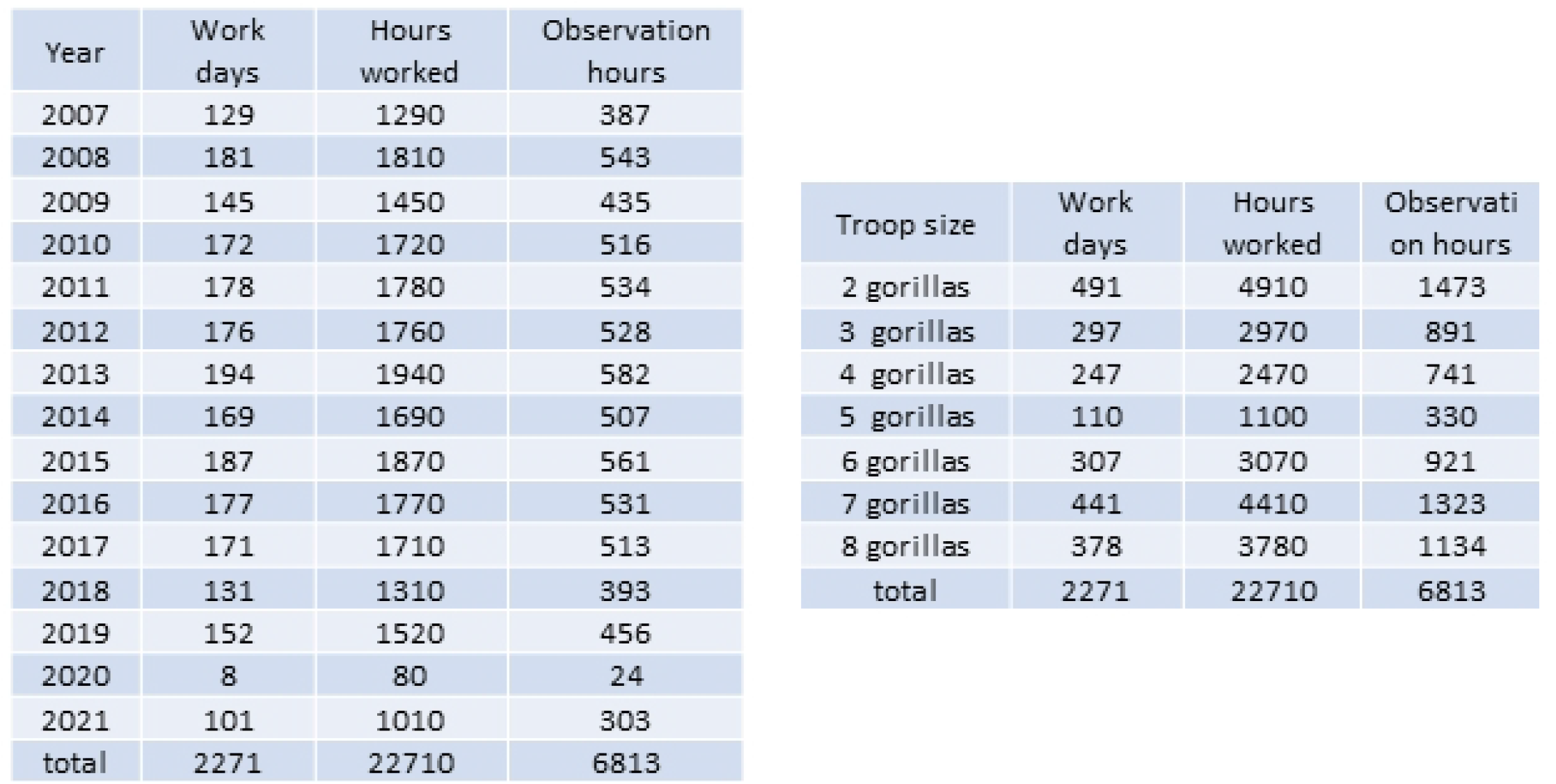
Distribution of observation time of gorillas.

**Fig. F2:**
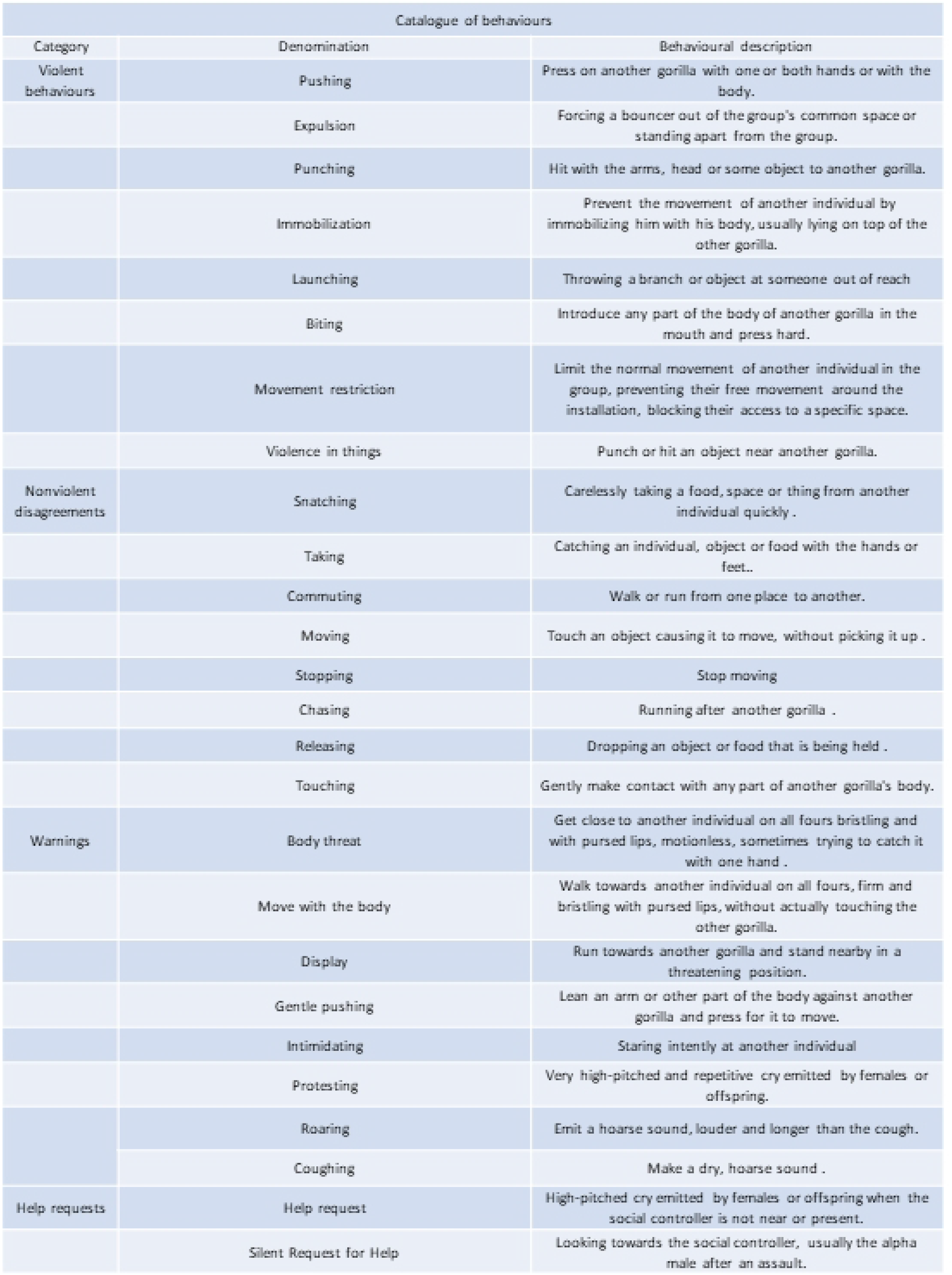
Catalogue of behaviours.

**Fig. F3.**
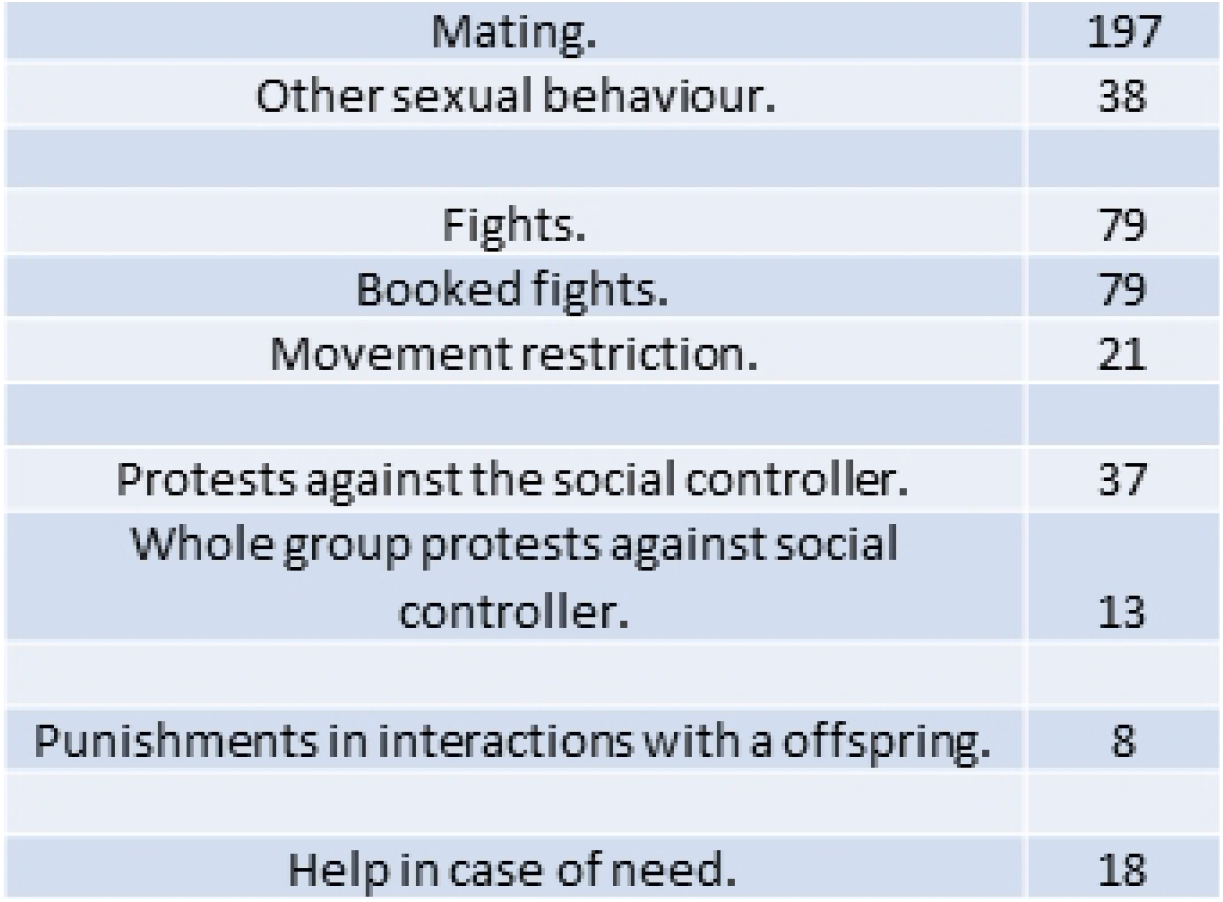
Remarkable behaviors observed in *ad libitum* sessions.

**Fig. F4.**
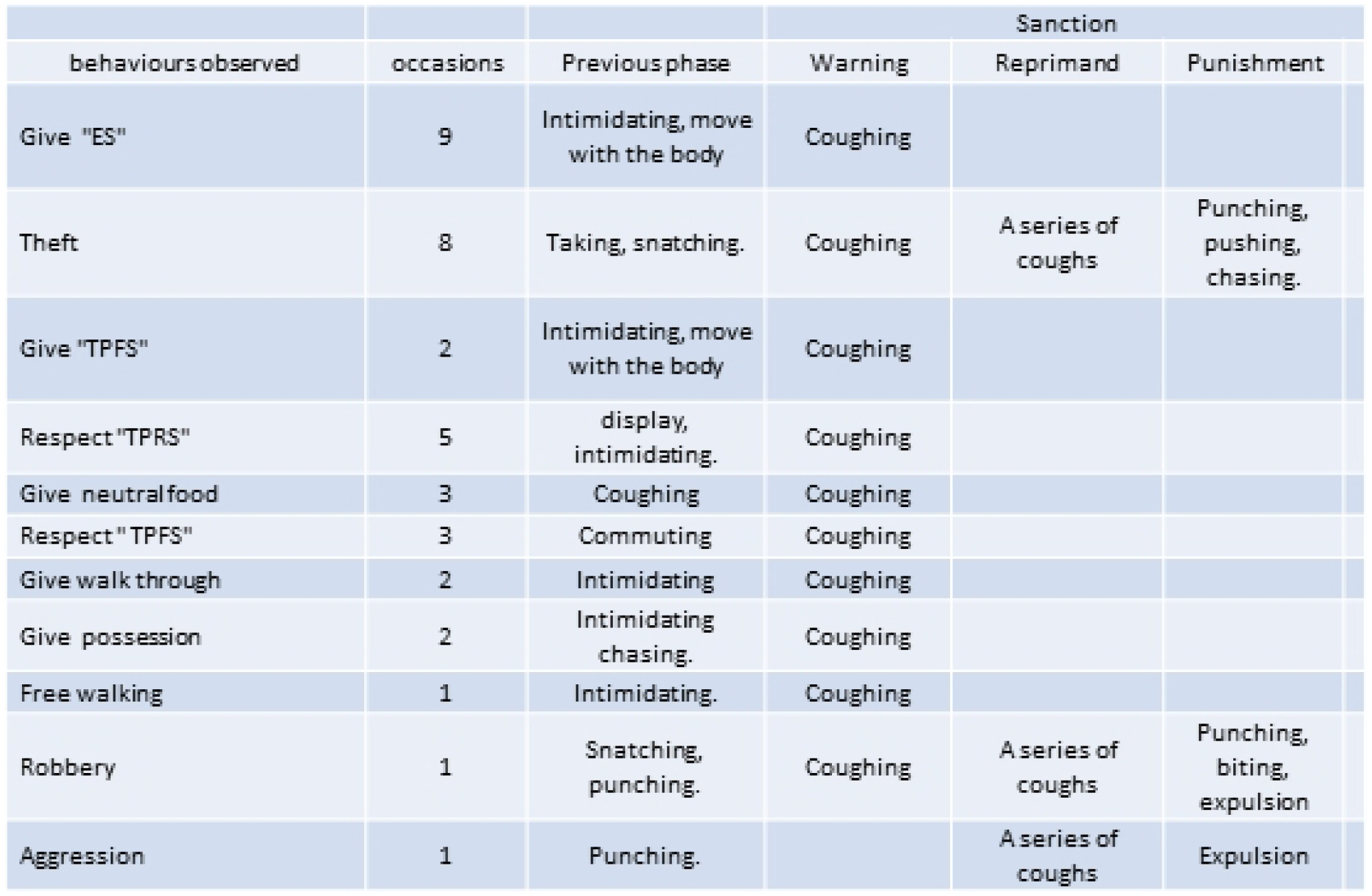
Behaviors observed in the phase of systematic observations through ethograms, which include the previous phase and the sanctioning phase.

**Fig. F5:**
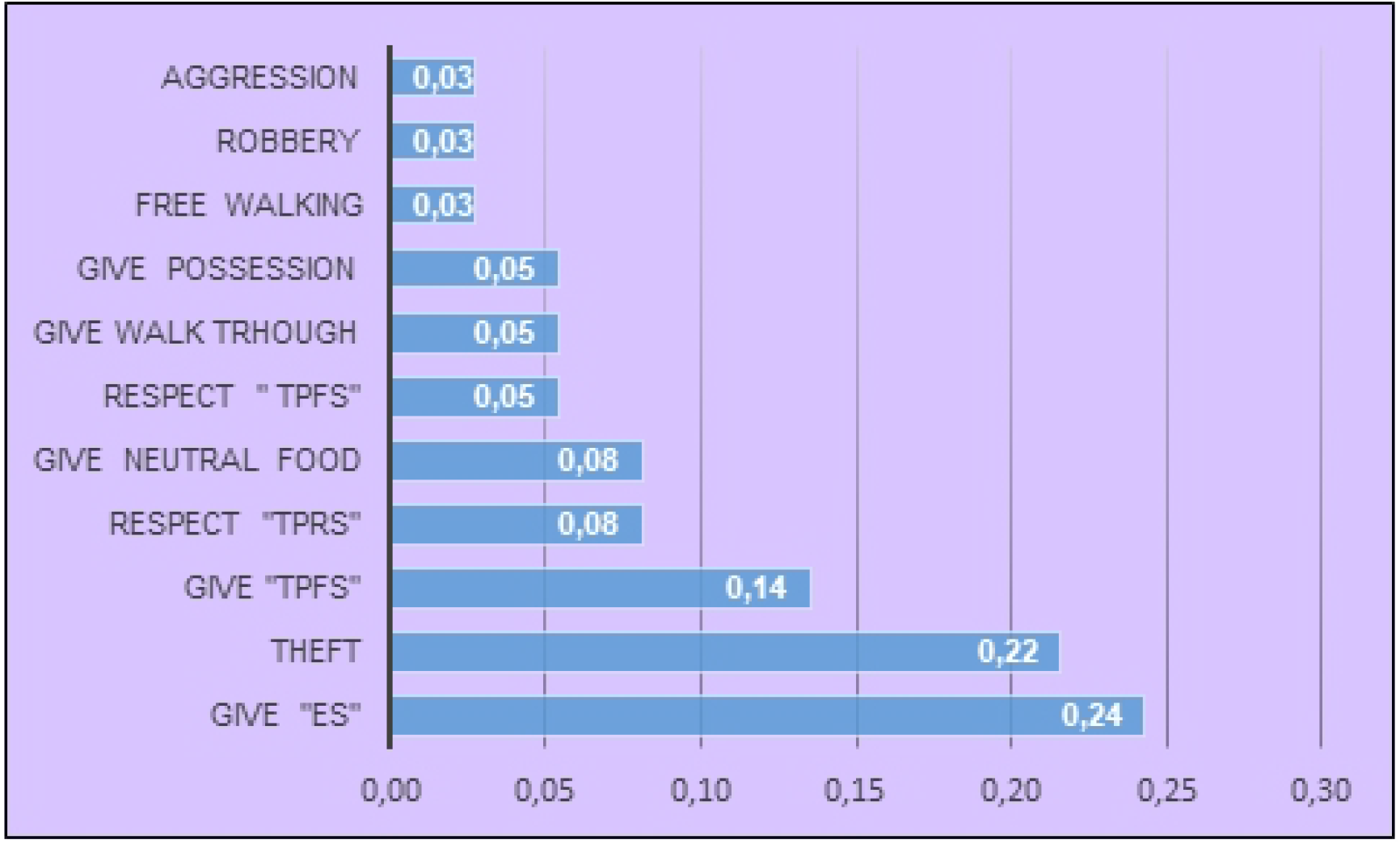
Frequency of norms obtained from observation using ethograms.

**Fig. F6:**
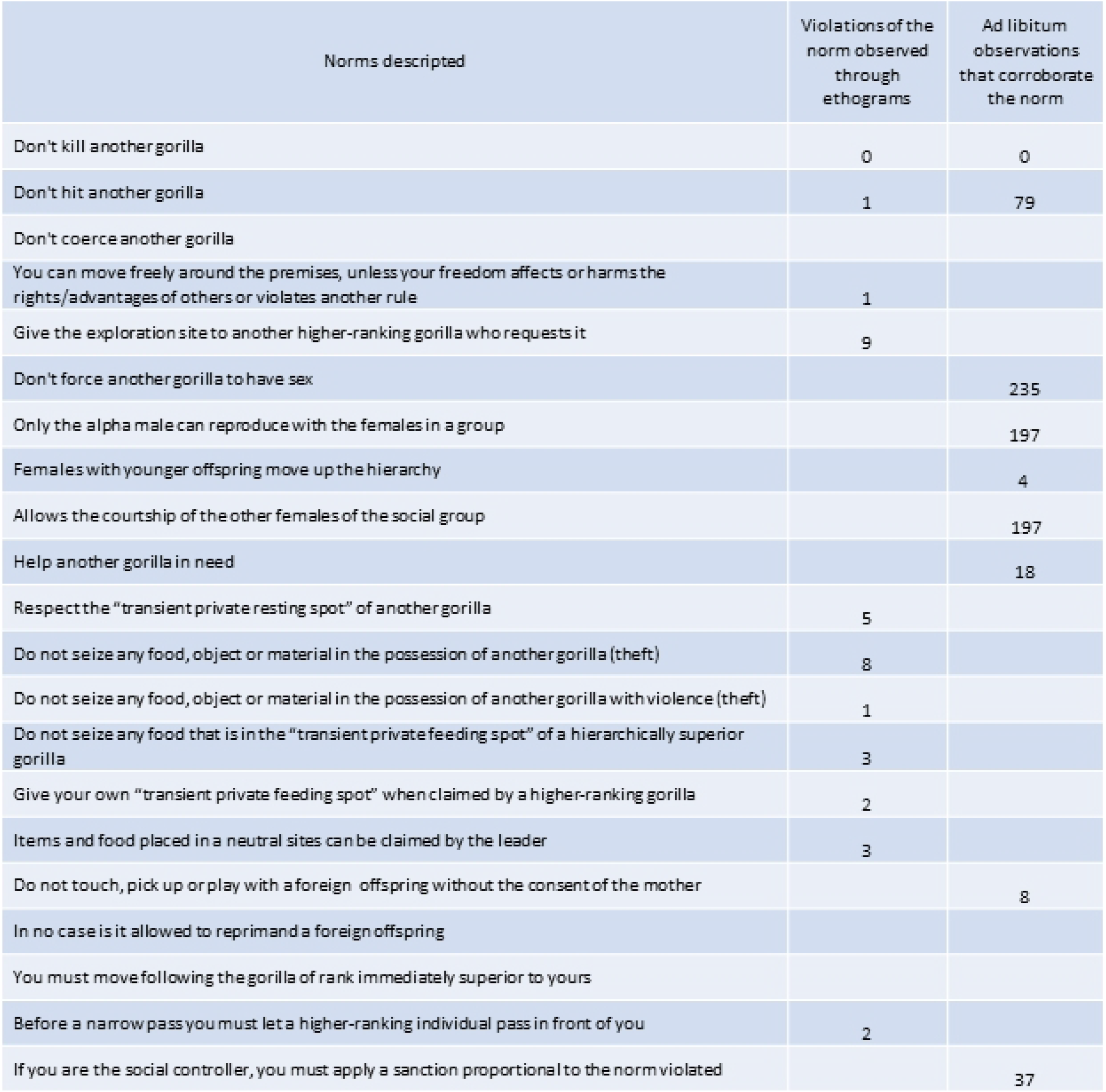
Summary of Recorded Norms. As expected, we did not find as much variety of norms at the delimited observation times as we did dur ng the ad lib tum observations. This is due to the fact that the systematic observation time is more limited, and that it is delimited to spec fic moments, it is not a continuous observation throughout the whole day like the observations dur ng the *ad libitum* phase. So the behavioural records are developed at times when only gorillas are being observed and no notable conflict may arise at that t me. In any case, we observe some regulations, the most common, such as transfers of food and space, or theft, which, as we see in the tab e, take p ace relatively frequently.

**Fig. F7:**
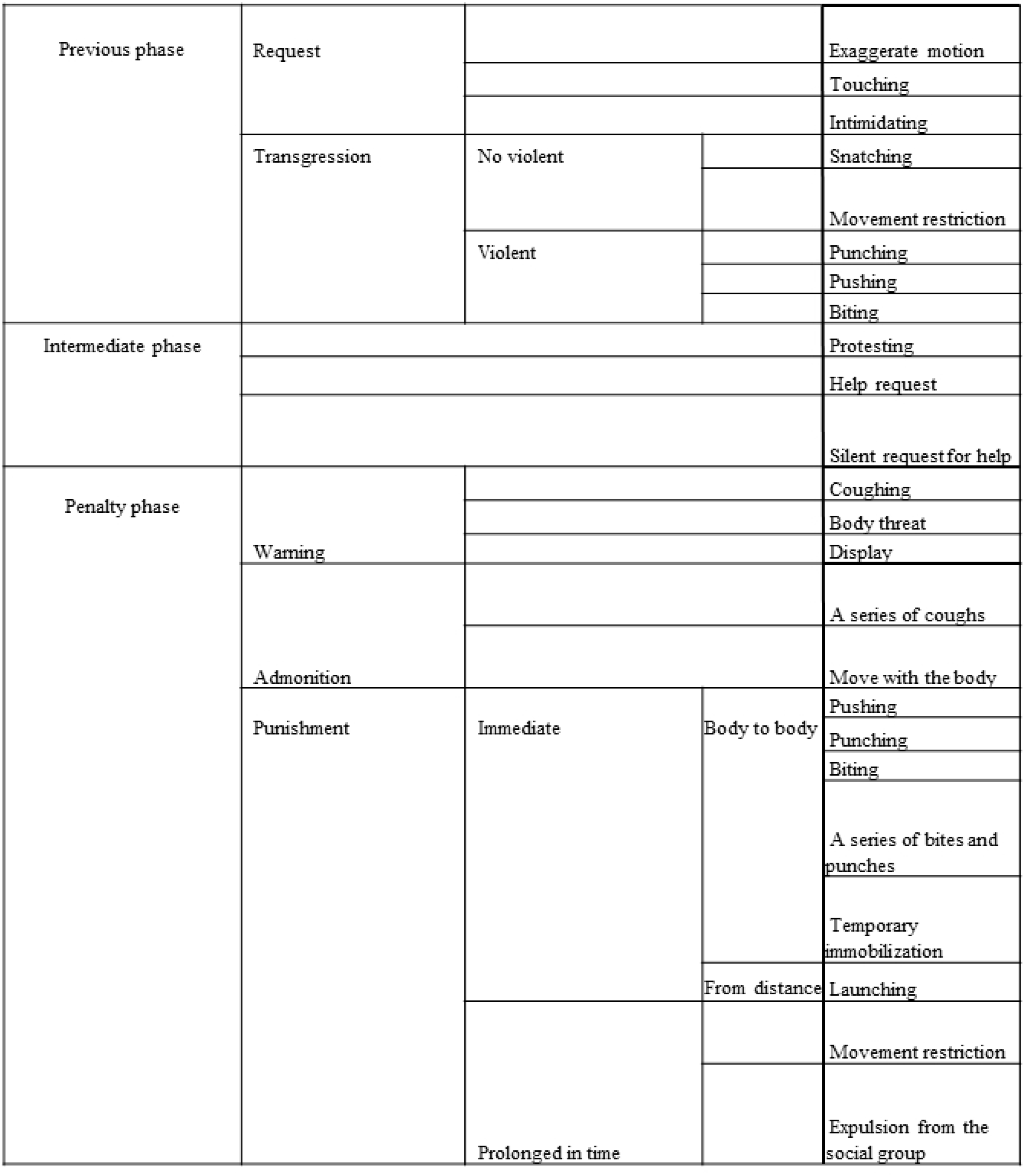
Phases in the. development of no1matire. behaviours ordered from lowest to highest severity.

## Materials and Methods

The study of the behaviour of gorillas requires many hours of observation, which makes it difficult to study them in their natural habitat, such as tropical forests with difficult access. For this reason, studies carried out in zoological institutions and sanctuaries that guarantee the maintenance of family structures while allowing prolonged observations and in different circumstances have gained interest. This research has been carried out from 2007 to 2021 in the gorilla facility of the Cabárceno Nature Park (Obregón, Spain). This group of gorillas live in captivity, which allows continued observation and investigation of their behaviour. The keeper team adopts care and management measures that allow the gorillas to develop their natural behaviours. Feeding and daily management follow the recommendations of the European Association of Zoos and Aquaria, always with respect to their needs.

Our research has been developed through *ad libitum* observations carried out in the afore mentioned period (F1). Regarding the work methodology, once the behaviours have been registered and catalogued, we have analysed their phenomenology and external characteristics, the circumstances that generated them and the reaction of other individuals. For this we have used the method of legal science, analysing the *facta* (the facts) to try to enunciate the *ius* (the rules that govern the expected behaviour). In order to systematize the results of the observations, we have taken as a point of reference, broadly speaking, the crimes contained in the Spanish Penal Code against personal legal interests, ranking them according to their importance (this is the criteria followed by the Spanish Penal Code). Depending on the protected legal interest, the Spanish Penal Code maintains the following order: homicides, injuries, attacks against freedom -coercion, illegal detentions-, sexual assaults, breaking and entering and theft and robbery.

In the hierarchy of behaviours, we have been guided by this criteria, but more complex criminal modalities have been ruled out in this investigation, first, because no indications have been detected that invite us to delve into them and, second, due to the difference in complexity between human social groups and gorilla social groups. Together with the above, we have introduced some more specific normative behaviours of the model of its social structure that, although, saving greater distances, could also have their equivalent in human societies (powers of the mothers, rules of hierarchies and rules for the mobility of the group). In the hierarchy of the catalogue of norms and values, we have also taken into account the seriousness of the sanction that the infraction deserves, so that we have considered that the greater the sanction, the more serious the infraction is considered and the more the underlying value is protected.

In the first moment of our research, the *ad libitum* observations allowed us to elaborate a catalogue of behaviours (F2), select what we have called “type-situations” (F6) and define both the normative behaviours and what we have designated as phases of the social response: previous phase, intermediate phase and sanctioning phase (F7).

Subsequently, we have carried out a smaller systematic study where we recorded behaviours during 21 hours of observation using ethograms. For this, we have used a multifocal sampling segmented by behavioural criteria focused on those behaviours that determine the normative behaviour of the individual. These systematic observations have taken place during variable time intervals between 30 minutes and 2 hours per session, where we have registered 37 normative behaviours described in table F4. The observations have been made at different times of the day, except for the night period, in order to check how they act depending on the circumstances (because their behaviour is not identical when they play and when they feed or during their rest period). The objective of this second phase of the study is to determine if the normative behaviours develop as we observed during the *ad libitum* phase and to corroborate that, indeed, we can verify a parallelism with the structure of the norms that govern human behaviour by prohibiting and sanctioning crimes (legal right, primary rule and secondary rule)3.

It is necessary to point out that many of the behaviours detected are occasional -because there are behaviours that are exceptions to the norm- hence we have to combine the two types of analysis, *ad libitum* and systematic. In the controlled observation sessions, which take place for a limited time, unusual behaviours do not always appear. However, during *ad libitum* observations, which are less regulated but more extensive, we have been able to observe them (F3). Obviously, in no instances have the behaviours been conditioned, provoked or claimed by the keepers or researchers, but rather they are completely spontaneous.

At the theoretical level, we have started by way of explaining the criminal phenomenon typical of Western societies with continental legal systems (dogmatic concept of crime) and from traditional theories about the purposes of punishment (retribution, general prevention and special prevention), but also using the contributions of criminology to the understanding of the criminal phenomenon. In any case, the concepts and terms used are in common use and their meaning is sufficiently consolidated in comparative criminal legal science.

Based on the observations made throughout the *ad libitum* phase, we have described the observed behaviours stating the regularization and the underlying rule, as well as the consequences derived from non-compliance and the exceptions detected. Once the two variables (behaviour and consequence) have been identified, we carry out a functional analysis –comparative between the normative system that we have described in our troop of gorillas and the one generally accepted in relation to human societies–. This is to determine if the relationship between behaviour and sanction observed can be functionally equivalent to the difference between a behaviour defined as a crime in a penal precept and its corresponding penalty. This process has two moments that are interrelated. Thus, once the expected behaviour (normative behaviour) has been stated, we apply a logical process of deduction to deduce the norm of conduct and the underlying value from this normative behaviour. This logical process of deduction is similar to the one used by the Criminal Doctrine to deduce from the criminal types (secondary norm) the normative mandate contained in the primary norm and the value that the criminal precept tries to protect (protected legal right). Next, we must verify that the infraction of the accepted norm (mandate or prohibition) generates a social response, by way of sanction, and, once this regularity has been verified, we try to state what the sanction consists of, what circumstances affect the seriousness of the sanction and in which cases the sanction is not imposed, despite meeting all the requirements for it. In all these cases, we are describing complex communicative social interrelationships in which three subjects participate: the transgressor (active subject), the injured party or victim (passive subject) and the social controller. In some interactions we have been able to observe the participation, in addition, of a *corbán* or propitiatory victim, that is a subject outside the conflict who suffers the violence used by the social controller to sanction an offending behaviour carried out by another subject.

At this point, and to describe, designate and define the observed phenomena, we have resorted to the contributions of penal doctrine, using its own terms and concepts as a point of reference. In this way we have identified normative behaviours (equivalent to criminal types); an underlying value (power, privilege or advantage equivalent to the protected legal asset); causes that allow the non-normative behaviour to be carried out (equivalent to the causes of justification, justification right of self-defence, legal duty or consent) or that prevent the imposition of a sentence (equivalent to being a minor).

From this sequence we deduce that, when a gorilla adjusts its behaviour to the normative mandate, it is reinforcing the validity of the norm and when a gorilla breaks any norm, it must be sanctioned. The regularity between non-compliance and sanction highlights the mandatory nature of the rule. Thus, we can conclude that some of the behavioural norms of gorillas fulfil the same functions as human legal norms, since they regulate behaviour patterns, are mandatory and are socially imposed. By observing these sequences of behaviours, we can deduce the ideal normative behaviour that each individual within the social group should exhibit. These normative behaviours may depend on the social hierarchy of each individual or be mandatory for all members of the group. We assume that behaviours that do not generate a sanctioning response or are not important for group interactions or are being carried out appropriately with respect to the norm. Therefore, when we observe that a gorilla –the active subject– carries out a behaviour to which another gorilla –the passive subject– responds with a complaint and/or leads to a reaction from the social controller, we can deduce that the active subject has carried out a behaviour contrary to what is expected (and required). From there, we can extract what the expected behaviour is, after comparing the behaviour actually produced with others observed in similar situations that did not generate a reaction or complaint.

## Results

We denominate “situation-type” those situations that generate expectations of behaviour. We have detected 21 type-situations (F6) in our research, so that the individual who is before them must act in a certain sense and, if he does not, will be sanctioned. To these behaviours that are subject to the norm that emerges from the situation-type, we will denominate “normative behaviours”. Behavioural expectations condition individuals’ behaviour in a similar way as legal norms do, so we deduce that, implicitly, such behaviours respond to norms. These norms are aimed at maintaining social order and among them we can distinguish: norms that define the hierarchy, rules that describe enforceable behaviours, rules that limit the power of the social controller, the silverback, and rules that regulate the sanctioning capacity and sanction. The social response to the behaviours that infringe the norm before the situation-type consist of three phases: a previous phase, in which before a situation-type the norm is transgressed, an intermediate phase, not essential, in which the subject who suffers the infraction claims or protests before the breach of a norm. In this case, we can find two possibilities, that the active subject complies with the rule or does not, in which case, the claim activates the social response to the infraction (and, therefore, becomes a kind of complaint), and a sanctioning phase, in which the social reaction to the infraction occurs (legal consequence) (F7).

The norms that are generated in the type-situations that we enunciate have the characteristics of reiteration of use, generality, claim of validity, stability and enforceability of the customary norms that, although they are common throughout history, are still maintained in our technological societies.

## Discussion: Valuation system and normative system

Human legal systems respond to a previous value system. This previous hierarchy of values can also be found in the norms that emanate from the observed type-situations, as well as in the social response to infractions. The values that underlie the norms of behaviour detected can be a power, a privilege or an advantage. They serve, not only to guide the correct behavior, but also to determine the modality and severity of the sanction that entails the breach of the norm.

Such rules may consist of mandates –which oblige one to act in a certain way– or prohibitions – which require refraining from acting–. We also find permits that authorize to infringe a mandate or a prohibition in a certain case. Most of the permits detected are related to powers granted to mothers over children or derived from the power of social control of the silverback. Sanctions are also organized according to their severity and are applied in proportion to the offence committed. Thus, we have made a first design of the regulatory system of the gorillas of Cabárceno, structured according to power, right, privilege or underlying advantage.

### Prohibitions and mandates

#### Protection of life

We can observe the absence of physical force to cause death. In this sense, life is a value, the most important and untouchable.

#### Protection of physical integrity and prohibition of aggression

In a second evaluation, physical integrity would also be protected, in this case, by prohibiting the use of physical force against another individual. We will consider “aggression” as any physical attack or use of physical force of one gorilla on another with the hand, foot, mouth or using an object as a weapon. Previous studies show that aggression is not common among western lowland gorillas^29,30^ and neither is it common in our group. However, when a gorilla (active subject) assaults another (phase 1), the latter (passive subject) can respond with “request for help” (phase 2) or also violently (“punching”, “pushing” or “biting”). In both cases, the social controller imposes a penalty (phase 3), even if no injury has occurred (F7). Fights are dissolved by the social controller and the individuals involved are reprimanded, even if no injuries have occurred.

The sanction is a necessary consequence of the aggression, but we have detected three cases in which the sanction is not imposed: self-defense in the face of imminent aggression; the use of force by the social controller in the exercise of his power of social control and, finally, in the case of offspring up to approximately 3 years old.

As for the sanction to be imposed, its severity depends on the value judgment that the infringing conduct deserves (judgment of action) or the effects of it (judgment of result). These criteria are valid in any case, so that, applying it to aggressions, they allow the sanction to be graduated as follows: depending on the judgment of action^31^, the penalty must be proportionate to the dangerousness of the conduct (F7). On the other hand, depending on the judgment of result^31^, the sanction depends on the intensity of the “request for help” –the more intense, the more serious the sanction– and on whether the aggression results in a fight or on whether the subject is a baby –due to his greater vulnerability–.

#### Protection of freedom

Freedom is also protected from coercion and movement restrictions imposed by third parties and constitute the third group of punishable offending behaviors. Coercion affects the “freedom to act” and occurs when an individual is forced to do something or not do something through the use of physical force. On the other hand, freedom of movement can be understood as the possibility that an individual has to determine by himself his situation in physical space^32^. Both –freedom to act and freedom of movement– deserve protection, but, unlike the assumptions analyzed previously, here the protection depends on the rank, so that each gorilla can move freely around the enclosure, unless it affects or harms the advantages of others or breaks a rule. Thus, the subordinate gorillas must give the “exploration spot” (ES) -the space through which a gorilla investigates in search of food and objects within the enclosureto the hierarchically superior gorillas who request it and to the silverback, who has absolute freedom. The regulation of freedom based on the hierarchy is complex, but it is interesting to note that, also in this case, the severity of the sanction depends, on the one hand, on the result produced and, on the other, on the nature of the “complaint” –intermediate phase–. The coercions or restrictions that occur in the context of play are exempt from sanctions although, in the event that the games take place with offspring, the freedom of the offspring to participate in it or the risk that this entails for them, is also taken into consideration to modulate the severity of the sanction. In any case, they are exempt from liability and, therefore, no penalty is imposed on the offspring.

#### Sexual self-determination and the use of violence in sexual relations

sexual relations for reproductive purposes are mediated by females^33^ in periods of ovulation, although sexual behaviors other than copulations between different members of the social group can also be observed; occasionally, even homosexual ones^34^. We have not observed that any gorilla –regardless of age, status or sex– has been forced to have sex or has tried to force another member of the group.

#### Protection of the rest space

We will call *transient private rest spot* (TPRS) the place of variable perimeter that each individual chooses to rest at a certain time. The TPRS of others is usually respected, from which we deduce that it could be assimilated to “their dwelling”^35^. Thus, it is forbidden to disturb a gorilla who is resting. Infringements of the rules governing the TPRS are subject to the criteria for exempting or aggravating the penalty described in relation to the protection of freedom.

#### Protection or respect of possession

Singularly complex is the normative framework that regulates and protects possession. Possession has already been found to be protected in other non-human primate species (7). In our study group, we have found two distinct modes of possession. The *physical possession* that is held over an object or food with which physical contact is established and as long as such contact is maintained^36^. And the possession of food found in the *transient private feeding spot* (TPFS), which comprises a space of variable perimeter around an individual containing food. The protection of physical possession does not depend on hierarchy, so it is forbidden for all members of the group to seize food or objects that others possess, regardless of their rank. However, the possession of food in the TPFS does depend on the status, so individuals of lower rank must give up their TPFS with the food it contains at the request of a higher-ranking gorilla. The hierarchy also grants advantages over objects or foods that are in a “neutral site” (NS), that is, those not under the possession of any gorilla. In this case, the social controller can claim them for himself remotely.

As for the sanction, the severity depends on the use or not of violence. The seizure without violence (theft) deserves less sanction than when it occurs with violence (robbery). Obviously, thefts are more numerous than robberies. The social controller is exempt from punishment although, in the rare occasion that an infringement occurs, the passive subject will usually protest. Also exempt from punishment are mothers with respect to what is possessed by the offspring and the offspring themselves.

#### Faculties of the mother over the progeny and the advantages of the offspring

the offspring enjoy a privileged status in which their safety within the group is guaranteed. On the other hand, every mother has recognized power over her offspring (parental authority) and they are the only ones who can sanction them (until weaning). All other members of the group are prohibited from touching, catching, playing or reprimanding an infant without its mother’s permission. In addition, in cases where a member of the group assaults an infant, the mother can temporarily acquire the role of the social controller and sanction them. When the social controller needs to impose a sanction, the severity will depend on the nature of the complaint or whether a danger has been created for the offspring, even if it occurs in the context of game.

#### Rules concerning the movement of the group

there are also rules that define the order in which gorillas must move in a group^37^, which establish what place each gorilla should occupy during joint movements, but this has already been the subject of previous research.

#### Functions of the social controller

The silverback is the leader of the group, a position that is accompanied by a role that includes privileges, powers and obligations. In the exercise of its role, it fulfils functions of social control, is the guarantor of the stability of the group and compliance with the rules and imposes sanctions, if appropriate, on those who infringe them. Their mere presence is a deterrent, avoids conflict, and allows for safer and longer-lasting social interactions (policing)^14^. In addition, it ensures compliance with the rules even against its own interests^38^. The sanctioning power it exercises, however, is not arbitrary, but it is subject to limits, so that if the sanction imposed is disproportionate or does not derive from an infraction, the females react by protesting or even persecuting it. In other words, the normative system of gorillas also contains norms that limit the exercise of social power to sanction, a function that, in human societies, is attributed to criminal law.

### Sanctions system

We have already stated that the infringement of the norm that is generated in a situation-type can lead to a sanction and that the social body that judges and sanctions is the silverback (social controller) and that the imposition of the sanction is modulated according to certain criteria (permits, exemptions, aggravations) and the initiative of the subject (complaint). The complaint, precisely, fulfils a double function: it sets in motion the “sanctioning process” and it determines the severity of the sanction according to the intensity of the complaint.

As for the criteria that govern the imposition of the sanction we can distinguish: circumstances that exempt from the sanction, which in turn can be objective (self-defense) or subjective (minor -offspring) and circumstances that aggravate the sanction, depending on the judgment of devaluing that the conduct deserves (Handlungs – und Erfolgsunwert). Thus, the use of violence^39^ or the dangerousness of the means used, deserve worse judgment of action and consequently, a more serious sanction.

Depending on the judgment of result, the infractions suffered by the offspring, or when they lead to a fight, are more punishable. However, we have not observed that the production of a more harmful result necessarily entails a greater sanction –unlike what usually happens in human legal systems^31^-. The severity of the sanction also depends on the existence, if any, and the intensity of the complaint.

The penalties are, depending on their severity (from less to more): “pushing”, “chasing”, “punching”, “biting”, “a series of bites and punches”, “temporary immobilization”, “restriction of movement” or “expulsion” (F2).

It is worth highlighting the parallels between the structure of the system of demanding criminal responsibility in human legal systems and that described in the troop of gorillas analyzed: society has rules that, depending on values (advantages, powers…) enunciate (situations-type) some guidelines of conduct (rules), the infringement of which generates a sanction, which is proportional to the way in which the infringement is carried out and the damage that has been caused. The complaint is not necessary when the silverback is the passive subject or when it is a fight, but, when it occurs, it initiates the sanctioning action. Even so, in some cases (exemptions), the penalty is not imposed^40^, either because the infringing conduct is allowed (causes of justification: self-defense or exercise of powers), or because the acting subject is not required to adapt his behavior to the normative mandate (exclusion of culpability: minors -offspring-).

## Conclusions

The western lowland gorillas troop that we have studied has a normative system that could even be qualified as “proto-normative system”, which breaks with the paradigm of human exceptionality in social systems. This system structures the group by coercively imposing norms (mandates and prohibitions) that regulate individual behavior in order to maintain social peace. To do this, it uses the threat of sanction (negative general prevention), sanctions proportionate to the infraction committed (retribution) and acts on individuals so that they do not commit more infractions in the future (special prevention). We have also found that there are exceptions to the imposition of the penalty similar to what would be our causes of justification and our causes of exclusion from culpability and circumstances that qualify it. In addition, the proto-system described contains rules that regulate the exercise of the powers of social control and limit the social reaction to crime, giving the members of the group the ability to react to arbitrariness and to limit the sanctioning power of the social controller.

This systematic set of rules and sanctions works as the criminal-legal system would. It is a system because the norms are coordinated with each other and has internal operating rules. And it is a normative system because the guidelines of behaviour that must be known and obeyed by the individuals to whom they are addressed and all individuals are subject to them. This similar regulatory system sets out punishable conduct, crimes and sanctions (penalties) that correspond to the social controller, who exercises functions of the State in human society and whose power is limited by the regulatory system itself, as also happens in human legal systems.

Limiting the research to a single group of gorillas will require corroborating our conclusions with larger studies, through ethograms that would give comparability to the research, to determine if they can be extended to the *Gorilla gorilla* species as a whole. But, in any case, it can serve as proof of the need to put an end to old presumptions in order to promote a new paradigm: that of the interspecific continuity of normative systems.

## Acknowledgments

We would like to thank Esther Hava García, Miquel Llorente Espino, Fernanda Genre, Isabel Esteban Gandarillas, Joel Clarke and the workers of the Cabárceno Nature Park for their collaboration. The investigation has complied with the ethical standards in the treatment of their animals with EC Guide for animal experiments.

## Author contributions

Conceptualization: PMCA, LGC

Methodology: PMCA, LGC

Investigation: LGC

Project administration: PMCA

Supervision: PMCA

Writing – original draft: PMCA, LGC

Writing – review & editing: PMCA, LGC

## Competing interests

Authors declare that they have no competing interests.

## Notes

### Competing Interest Statement

The authors have declared no competing interest.

## References and Notes

1. H. Kelsen, Teoría General de Las Normas, (Trillas, México, 1994).

2. M. Atienza Rodriguez, J. RUÍZ Manero, Las piezas del derecho, (GSB, Barcelona, 2007).

3. F. Gudin RodrÍguez-MagariÑos, Ubi societas, ibi ius: sobre las normas que organizan a los animales gregarios. Ius et scientia, 2 (1), 1–34 (2016).

4. E. Durkheim, Las reglas del método sociológico, (Ernestina de Champourun trad., Fondo de cultura económica, México, ed. 2, 2001).

5. D. J. Melnick, The genetic consequences of primate social organization: a review of macaques, baboons and velvet monkeys. Genetica, 73 (1987).

6. M. Bekoff, Wild justice and fair play: cooperation, forgiveness, and morality in animals. Biology and Philosophy, 19 (2004).

7. H. Kummer, Primate societies: group techniques of ecological adaptation. (W. Goldschmidt ed. Aldine Atherton inc, Chicago, 1971).

8. S. Yamamoto, A. Takimoto, Empathy and Fairness: Psychological Mechanisms for Eliciting and Maintaining Prosociality and Cooperation in Primates. Soc. Just. Res., 25 (2012).

9. S. F. Brosnan, R. Bshary, Cooperation and deception: from evolution to mechanisms. Philosophical Transactions of the Royal Society B: Biological Sciences, 365,1553 (2010).

10. S. F. Brosnan, Justice-and fairness-related behaviors in nonhuman primates. Proceedings of the National Academy of Sciences, 110, Supplement 2, 10416–10423 (2013).

11. J. Pierce, M. Bekoff, Wild justice redux: What we know about social justice in animals and why it matters. Social Justice Research, 25(2) (2012).

12. F. B. M. De Waal, Good Natured: the origins of right and wrong in humans and other animals (Harvard University Press, Cambridge, 1996).

13. H. Sigg, J. Falett, Experiments on respect of possession and property in hamadryas baboons (Papio hamadryas). Animal Behavior, 33 (1985).

14. J. C. Flack, F. B. M. De Waal, D. C. Krakauer, Social structure, robustness, and policing cost in a cognitively sophisticated species. The american naturalist, vol. 165, nº 5 (2005).

15. E. B. Tylor, Primitive Culture: Researches into the Development of Mythology, Philosophy, Religion, Art, and Custom. (Ed. J. Murray, London, 1871).

16. J. Sabater Pi, El chimpancé y los orígenes de la cultura, (Anthropos, Barcelona, ed. 3, 1992).

17. R. Salmi, K. Hammerschmidt, D. M. Doran-Sheehy, Western Gorilla Vocal Repertoire and Contextual Use of Vocalizations. Ethology 119, nº 10 (2013).

18. D. Hedwig, R. Mundry, M. M. Robbins, C. Boesch, Contextual Correlates of Syntactic Variation in Mountain and Western Gorilla Close-Distance Vocalizations: Indications for Lexical or Phonological Syntax? Animal Cognition 18, º 2 (2015).

19. A. H. Harcourt, K. T. Stewart, Function and meaning of Wild Gorilla close calls. Behaviour 133, º 11 (1996).

20. E. Genty, T. Breuer, C. Hobaiter, R. Byrne, Gestural communication of the gorilla (Gorilla gorilla): Repertoire, intentionality and possible origins. Animal cognition, 12. 527–46 (2009).

21. M. Klailova, P. C. Lee, Wild Western Lowland Gorillas Signal Selectively Using Odor. PLOS ONE 9, º 7 (2014).

22. G. Cordoni, E. Palagi, S. Tarli, Reconciliation and Consolation in Captive Western Gorillas. International Journal of Primatology, 27 (2006).

23. E. J. C. Van Leeuwen, E. Zimmermann, M. DAVILA Ross, Responding to inequities: gorillas try to maintain their competitive advantage during play fights. Biology Letters, 7, p(2011).

24. C. Tephan, Third-Party Intervention in Wild Western Lowland Gorillas (Gorilla Gorilla Gorilla). International Journal of Primatology 40, nº 6 (2019).

25. Y. Iwata, Food dropping as a food transfer mechanism among western lowland gorillas in Moukalaba-Doudou National Park, Gabon. Primates, 55(3):353–8 (2014).

26. A. K. Kalan, H. J. Rainey, Hand-clapping as a communicative gesture by wild female swamp gorillas. Primates, 50 (3):273–5, 2009.

27. D. P. Watts, “Gorilla social relationships: a comparative overview”,in Gorilla biology: a multidisciplinary perspective, (Cambridge University Press, Cambridge, pp. 302–328, 2003).

28. M. M. Robbins, C. Ando, K. A. Fawcett, C. C. Grueter, D. Hedwig, Y. Iwata, J. L. Lodwick, S. Masi, R. Salmi, T. S. Stoinski, A. Todd, V. Vercellio, J. Yamagiwa, Behavioral variation in gorillas: evidence of potential cultural traits. PLOS ONE, 11(9): e0160483 (2016).

29. E. J. Stokes, Within-group social relationships among females and adult males in wild western lowland gorillas (Gorilla gorilla gorilla). American journal of primatology, 64 (2004).

30. T. L Maple, M. P. Hoff, Gorilla behavior. (Van Nostrand Reinhold Company, New York, 1982).

31. H. H. Jescheck, Tratado de Derecho Penal parte general, (Traducción y adiciones de Derecho español de S. Mir Puig y F. Muñoz Conde, ed. BOSCH Casa editorial S.A., Barcelona, 1981).

32. F. MuÑÓz Conde, Derecho Penal parte especial. (Tirant lo Blanch, Valencia, ed. 23, 2021).

33. T. S. Stoinski, B. M. Perdue, A. M. Legg, Sexual behavior in female western lowland gorillas (Gorilla gorilla gorilla): evidence for sexual competition. American Journal of Primatology, 71(7):587–93 (2009).

34. C. Grueter, T. S. Stoinski, Homosexual behavior in female mountain gorillas: reflection of dominance, affiliation, reconciliation or arousal? PLoS ONE 11(5):p e0154185 (2016)

35. P. M. De La Cuesta Aguado, “Allanamiento de morada, domicilio de personas jurídicas y establecimientos abiertos al público”, in Tratado de Derecho Penal parte especial (I): delitos contra las personas, (Tirant lo Blanch, Valencia, ed. 3, 2021).

36. L. Díez-Picazo, A. GullÓn, Sistema de Derecho Civil, vol. III, (tomo I), (Tecnos, Madrid, ed. 10, 2019).

37. G. B. Schaller, The year of the gorilla, (University of Chicago press, Chicago, 1988).

38. F. B. M. De Waal, Evolutionary ethics, aggression, and violence: lessons from primate research. Journal of Law, Medicine and ethics, 32 (2004).

39. S. Mir Puig, Derecho Penal parte general, (Reppertor, Barcelona, ed. 10, 2016).

40. P. M. De La Cuesta Aguado, Tipicidad e imputación objetiva, (Tirant Lo Blanch,Valencia,1996).

